# Integrated epigenome and transcriptome analysis of normal and arrested meiotic initiation during mouse spermatogenesis

**DOI:** 10.1101/2022.02.18.481086

**Authors:** Xiaoyu Zhang, Sumedha Gunewardena, Ning Wang

## Abstract

The transition from mitotic to meiotic cell cycles is a transcriptional event that entails the activation of genes important for meiosis and requires germline-specific retinoic acid (RA) signaling target, *Stra8*. To identify novel transcription factors underlying mammalian meiotic initiation, we conducted integrative snATAC-seq and scRNA-seq analysis using wild-type and *Stra8*-deficient mouse testicular cells to map the chromatin accessibility and gene expression landscapes of normal and genetically arrested meiotic initiation. Our results identified a cluster of putative inhibitory transcription factors for meiotic initiation, which we consider “meiotic inhibitors”. STRA8 binds to the regulatory sequences of these meiotic inhibitors and represses their expression upon meiotic initiation. In *Stra8*-deficient cells that suffer meiotic initiation arrest, the chromatin accessibility of these meiotic inhibitors is increased, concurrent with their uncontrolled and sustained expression. Among these meiotic inhibitors include KLF4, MAX, and MAZ. Importantly, by analyzing the single cell transcriptomes of human testes, our data show that these putative meiotic inhibitor genes are upregulated in early germ cells from patients with spermatogenic failure. Our study suggests that proper repression of meiotic inhibitors is essential for both mouse and human spermatogenesis.

## Main

Sexual reproduction depends on the production of haploid gametes through meiosis, a defining feature of germline cells. The transition from mitotic to meiotic cell cycles is a transcriptional event, which entails the activation of meiotic genes responsible for meiotic chromosomal events, including synapsis and recombination (*1, 2*). In budding yeasts, nutrient restriction of diploid cells induces their meiotic entry, and this process requires *inducer of meiosis 1* (*IME1*), which encodes a single master transcription factor inducing meiotic gene activation (*3*). Despite that many meiotic genes are conserved from yeasts to mammals, the molecular machinery responsible for their activation in mammalian germ cells remains elusive. *Stimulated by retinoic acid gene 8* (*Stra8*) downstream of retinoic acid (RA) signaling is considered as a best characterized gatekeeper of meiotic initiation. *Stra8*-deficient germ cells is required for meiotic gene activation *in vitro* and *in vivo* (*4, 5*). Although STRA8 has been demonstrated to bind to a broad spectrum of gene promoters, including meiotic genes (*6*), other studies suggest that STRA8 induces meiotic gene activation indirectly, e.g., the autophagy pathway (*7*). In support of this notion, neither STRA8 alone nor with its cofactor MEIOSIN, is sufficient to induce meiotic initiation (*8*). Recently, nutrient restriction has been shown to induce meiotic initiation in male germline stem cell culture, suggesting a conserved role for the nutrient restriction pathway in meiotic gene activation (*5*). Together, transcriptional regulation of meiotic initiation represents a major gap of knowledge that hinders our understanding of meiotic fate specification during mammalian germline development.

Assay for Transposase-Accessible Chromatin using sequencing (ATAC-seq) provides genome-wide profiles of open-chromatin regions or chromatin accessibility – a proxy for *cis*-regulatory element (CRE) activity (*9*). Through complex CRE interactions, transcription factors (TFs) act to modulate gene expression. To overcome the limited resolution of bulk-tissue ATAC-seq due to cell-type heterogeneity, single cell approaches have recently been adopted to study CRE activity in normal tissues that establish cellular identity or disease pathogenesis at the single cell resolution.

To profile the epigenomic and transcriptomic landscape of mammalian germ cell entry into meiosis, we conducted snATAC-seq (10X Genomics) and scRNA-seq (10X Genomics) using dissociated cells from contralateral testes of wild-type and *Stra8*-deficient mice at postnatal day 12 (P12) and P21, when early male germ cells are enriched (**fig. S1**). After quality-control and filtering (**fig. S2-S4**), 32,657 single-nucleus epigenomes and 22,839 single-cell transcriptomes were retained (**fig. S5 and S6**). Dimension reduction using uniform manifold approximation and projection (UMAP) followed by clustering analysis identified distinct cell-type specific clusters in the batch-corrected snATAC-seq and scRNA-seq (**fig. S5 and S6**). In snATAC-seq, based on chromatin accessibility at the promoter regions of known marker genes, we profiled major germ cell types, which include spermatogonia (Spg, 9,814 nuclei), spermatocyte (Spc, 3,451 nuclei) and spermatid (Std, 1,008 nuclei), as well as major somatic cell types, which include Sertoli cell (Stl, 4,293 nuclei), myoid cell (Myd, 4,256 nuclei) and Leydig cells (Lyd, 9,835 nuclei) (**fig. S5**). Transcriptome analysis detected similar phenotypes, including spermatogonia (Spg, 3,941 cells), spermatocyte (Spc, 4,924 cells), spermatid (Std, 710 cells), Sertoli cell (Stl, 9,488 cells), myoid cell (Myd, 3,531 cells) and Leydig cells (Lyd, 245 cells) (**fig. S6**). The distributions of different cell types in wild-type and *Stra8*-deficient testes at P12 and P21 are consistent in snATAC-seq and scRNA-seq analyses. To compare the consistency between cluster assignment in the snATAC-seq and the scRNA-seq datasets, we pooled each snATAC-seq cluster and derived gene activity score for the top 2,000 highly variable genes. Pearson’s correlation coefficient between each snATAC-seq cluster and scRNA-seq cluster indicated good concordance between the two datasets (**fig. S6E**).

To identify transcriptional program critical for meiotic initiation, we selected germ cell epigenomes (spg, spc, and std subsets) from WT and *Stra8*-deficient mice for further analysis. Pooled epigenomes from WT and *Stra8*-deficient samples identified 9 cell clusters (**Fig. 1A**). Based on known marker genes, we annotated these 9 clusters as male germ cells at consecutive differentiation stages from SSC to spermatid 2 (Std2) through meiotic prophase I (leptotene, zygotene, pachytene, and diplotene) (**Fig. 1B**). Notably, the epigenomes of *Stra8*-deficient germ cells are arrested at a leptotene-like stage, the first phase of meiosis prophase (**Fig. 1A, C**). We identified 183,122 total reproducible peaks in wild-type cells (**Fig. 1D**), of which 141,216 were classified as differentially accessible peaks across all clusters. In *Stra8*-deficient cells, we identified 151,023 total reproducible peaks (**Fig. 1E**). To describe the patterns of these peaks during spermatogenesis, we used an iterative clustering procedure. Most of the peaks are cell type-specific (**Fig. 1F, 1G**). Notably, clusters 7 and 8, which are mostly in germ cells at meiotic prophase I, are enriched for GO terms related to meiosis, suggesting that the regulatory sequences of genes important for meiosis become more accessible to allow for their expression during meiosis prophase (**Fig. 1F**). Although both wild-type and *Stra8-deficient* cells show similar CREs with GO function related RNA processing at leptotene stage, CREs important for meiosis are not accessible in *Stra8*-deficient germ cells (**Fig. 1G**). Thus, we consider that *Stra8*-deficient cells enter a “leptotene-like” stage.

**Figure 1.**
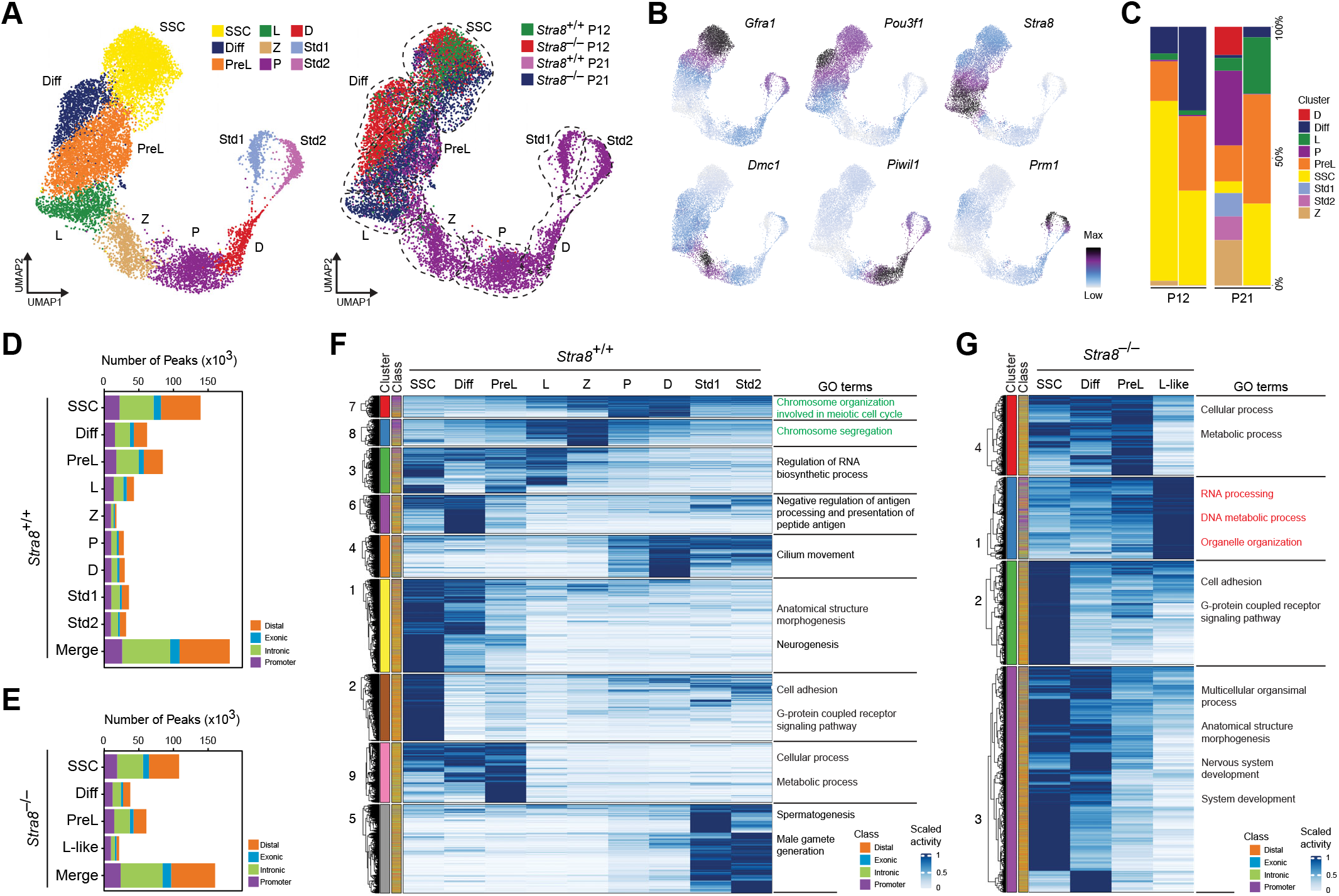
snATAC-seq analysis of WT and *Stra8*-deficient juvenile testes. (**A**) Uniform manifold approximation and projection (UMAP) plot of germ cells captured from WT and *Stra8*-deficient samples in P12 and P21 colored by cluster and sample identity. (**B**) Different stage germ cell marker expression shown as UMAP feature. (**C**) Histograms showing the percentage of each time point in each germ cell cluster. (**D-E**) Peak calls from each cluster in WT and *Stra8*-deficient are shown individually that merged peak set. Color indicates the type of genomic region overlapped by the peak. (**F-G**) Clusters of *cis*-regulatory element (CRE) activity across cell types in WT (**F**) and *Stra8*-deficient (**G**) testes. CREs are grouped by activity cluster (k-means followed by hierarchical clustering) and genomic class (left). Shown on the right are representative enrichments for biological processes.

We extracted leptotene epigenomes (320 from WT sample and 818 from Stra8-deficient sample) and conducted sub-clustering, which identified three clusters. While C2 cluster includes most WT cells, *Stra8-deficient* cells are mostly in C1 and C3 clusters (**Fig. 2A)**. By performing differential accessibility analysis, we observed 1,262 peaks (0.91%) with increased accessibility and 1,433 peaks (1.5%) with decreased accessibility in *Stra8-deficient* cells when compared to WT cells (**Fig. 2B**). Next, we conducted a global analysis of differential accessibility between WT and *Stra8-deficient* cells by calculating pseudobulk averages from snATAC-seq. We categorized differential accessibility as peaks enriched in WT cells and peaks enriched in *Stra8-deficient* cells (**Fig. 2C**). Quantitative color-coded enrichment plots of aggregated data from each category are shown in the top row (**Fig. 2C**). In *Stra8-deficient* cells, we found that, while 514 genes exhibit decreased chromatin accessibility, 1,044 genes exhibit increased chromatin accessibility near transcription start site (TSS), suggesting a role for STRA8 in reducing chromatin accessibility. To determine whether STRA8 binds to these regulatory sequences with differential chromatin accessibility, we analyzed a published ChIP-seq dataset (*6*). We found that STRA8 binds strongly to the genes with decreased accessibility in *Stra8-deficient* cells but weakly to the genes with increased accessibility (**Fig. 2D**). Notably, only peaks with higher chromatin accessibility in WT cells is annotated to genes enriched with GO related to meiosis (**Fig. 2E**). We then measured the deviation scores and variations of transcription factor (TF) (**Fig. 2F**). Interestingly, this analysis identified that TFs, such as Sohlh2, Ctcfl, Crem and Smad1, are enriched in WT cells, suggesting that STRA8 is required for their expression. Interestingly, Sohlh2 is required for spermatogonial differentiation (*10*). In contrast, Klf4, Maz, Tcf12 and Max are enriched in *Stra8-deficient* cells, suggesting that STRA8 represses their expression. Max is known to prevent meiotic fate in embryonic stem cells and male germ cells (*11*). To test the enrichment of specific genes, we computed the variability of these enrich TFs. Interestingly, *Klf4*, an essential pluripotency gene, is among the most significantly enriched variability TFs (**Fig. 2G**). Similarly, for *Stra8-deficient* cells that resulted in strong motif accessibility differences compared to WT cells, the motifs identified were consistent with the targeted transcription factor, as shown for *Klf4* (**Fig. 2H**).

**Figure 2.**
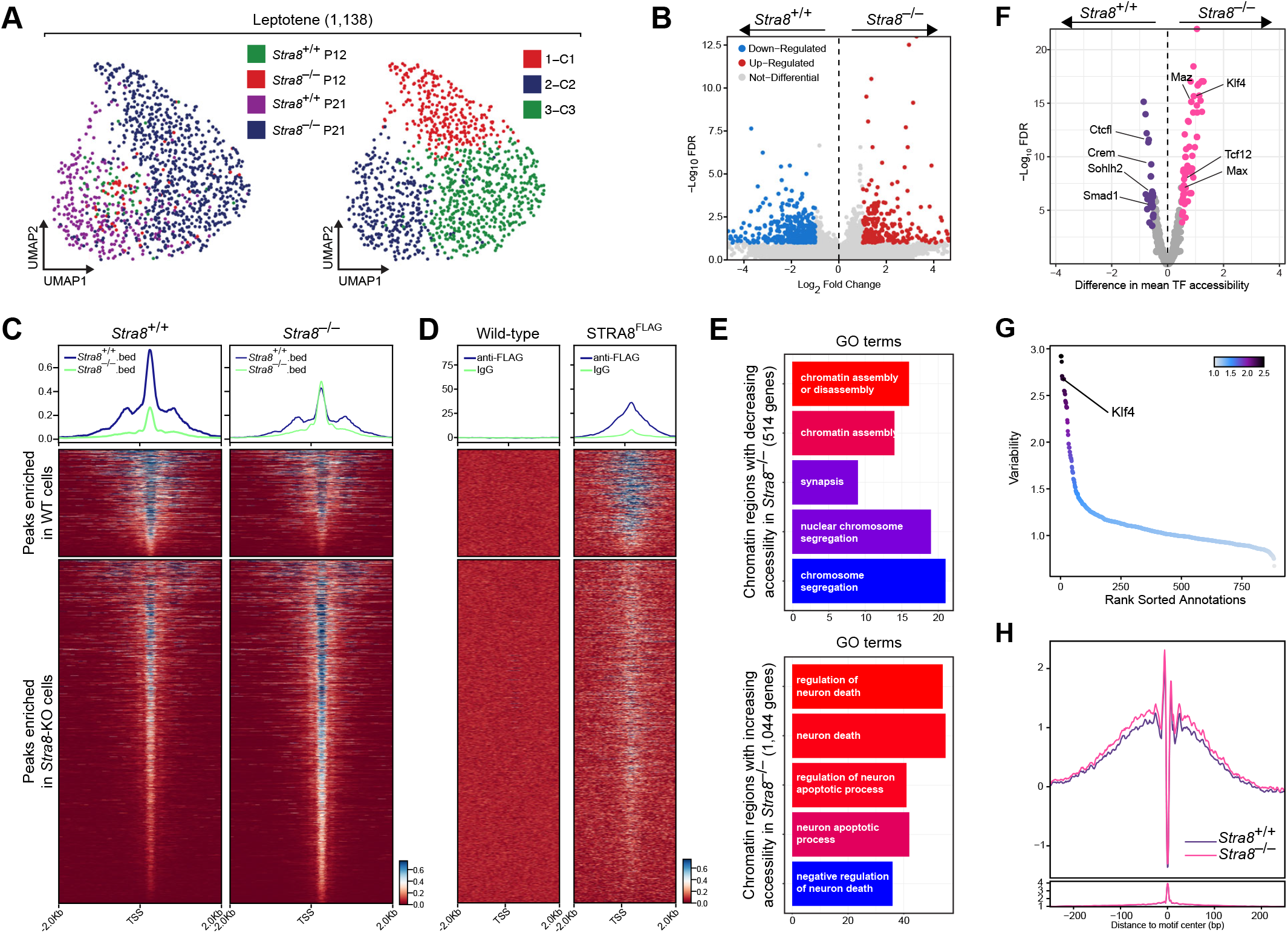
Loss of STRA8 leads to dysregulated chromatin accessibility in leptotene-like spermatocytes. (**A**) Uniform manifold approximation and projection (UMAP) plot of Leptotene stage cells captured from wildtype and Stra8-knockout in day 12 and 21 colored by cluster and sample identity. (**B**) Volcano plots displaying differential accessibility analysis results based on MACS2-defined peak regions for wild-type and *Stra8*-deficient from leptotene stage. (**C**) Identification of putative regulatory elements in wild-type and *Stra8*-deficient from leptotene stage. (**D**) ChIP-seq assay for STRA8 (Data analyzed from Kojima et al, Elife, 2019). (**E**) Top 5 significantly enriched GO terms sorted by GeneRatio identified on the basis of genes annotated to regions more accessible due to wild-type and Stra8-knockout are shown. (**F**) Volcano plots showing the differential TF motif accessibility using the mean TF. (**G**) Motif accessibility in the chromVAR TF bias-corrected deviation in wild-type and Stra8-knockout samples. (**H**) Visualization of variability transcription factor (TF) binding motif results for the specifically accessible regions in leptotene by using the TF motif database.

We compared the chromatin accessibility profiles in *Klf4* locus of WT and *Stra8-deficient* cells. The chromatin accessibility of *Klf4* locus exhibits a reversible decrease in WT samples during meiotic initiation, in that it is progressively decreased from SSC to meiotic prophase (zygotene), followed by an increase from zygotene to Std2 (**Fig. 3A**). In contract, the chromatin accessibility of KLF4 in *Stra8*-deficient sample is sustained till the leptotene-like stage. Based on the peaks, we computated the gene scores (**Fig. 3B**). Increased gene score of *Klf4* is strongly linked with *Klf4* expression, which is measured from scRNA-seq dataset. and decreased accessibility of genomic regions that KLF4 motifs compared with *Stra8*-deficient (**Fig. 3B**). Using multi-omic data from STRA8 ChIP-seq and RNAP ChIP-seq for nascent *Klf4* RNA transcription, we defined the interaction regions of STRA8 with *Klf4* (**Fig. 3C and D**). To confirm that *Klf4* is upregulated in *Stra8*-deficient cells, we conducted in-situ hybridization (ISH) for *Klf4* mRNA (**Fig. 3E**) and immunostaining for KLF4 protein (**Fig. 3F**). Thus, our data support the model that STRA8 represses *Klf4* expression during meiotic initiation in WT germ cell development; in *Stra8*-deficient germ cells that fail to enter meiosis, KLF4 expression is sustained due to loss of STRA8 (**Fig. 3G**).

**Figure 3.**
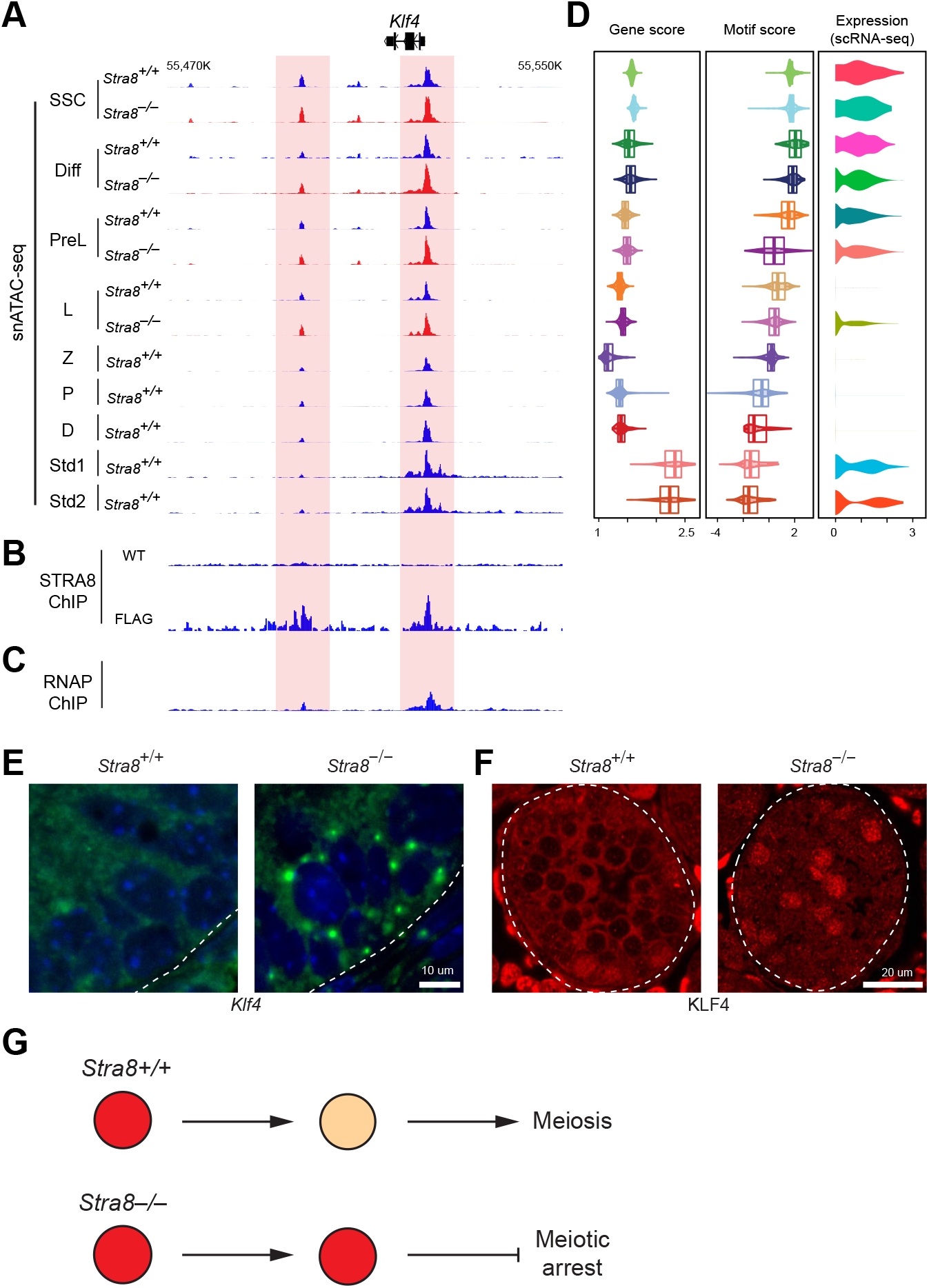
*Klf4* is a direct repressive target of STRA8. (**A**) *Klf4* genome accessibility tracks for each stage in WT and *Stra8-deficient* samples by snATAC-seq. (**B**) STRA8 ChIP-seq, which shows STRA8 binding to *Klf4* peaks (shaded areas). (**C**) RNAP ChIP-seq, which shows nascent transcription sites of *Klf4*. (**D**) Violin plots of gene score, motif score, and expression in WT and *Stra8-deficient* samples. ISH for *Klf4* mRNA (**E**) and immunofluorescent staining for KLF4 protein (**F**) in WT and *Stra8-deficient* testicular cross sections. (**G**) Schematic shows that *Stra8-deficient* differentiating male germ cells exhibit sustained KLF4 expression and suffer meiotic arrest.

In summary, our study establishes a new paradigm, in which the commitment to meiotic fate in germ cells requires STRA8-mediated repression of meiotic inhibitors. Furthermore, TF identification is challenging in scRNA-seq data, since the expression of several cell type-specific TFs is low. Here, our integrative snATAC-seq and scRNA-seq dataset addresses this challenge by allowing the identification of key regulatory logics for each testicular cell type through CRE, motif enrichment, and TF-target gene co-expression.

## Supporting information

Complete Supplementary Figures

## Acknowledgments

We thank the University of Kansas Medical Center (KUMC) Genomic Sequencing Facility for expertise in library preparation and sequencing.

## Funding

National Institutes of Health grant R01 HD103888 (NW)

KUMC Lied Pre-Clinical Research Pilot Grant Program (NW)

National Institutes of Health grant P20 GM103418 K-INBRE Postdoctoral Fellowship Award (XZ)

KUMC BRTP Postdoctoral Fellowship Award (XZ)

Department of Molecular and Integrative Physiology at KUMC (NW)

## Author contributions

Conceptualization: XZ, NW

Methodology:XZ, NW

Investigation: XZ, SG

Visualization: XZ, NW

Funding acquisition: NW

Supervision: NW

Writing – original draft: XZ, NW

Writing –review & editing: XZ, NW

## Competing interests

Authors declare that they have no competing interests.

## Notes

### Competing Interest Statement

The authors have declared no competing interest.

