## Supplementary material for "Integrated epigenome and transcriptome analysis of normal and arrested meiotic initiation during mouse spermatogenesis": Complete Supplementary Figures

**fig. S1.** Workflow of the experiments.

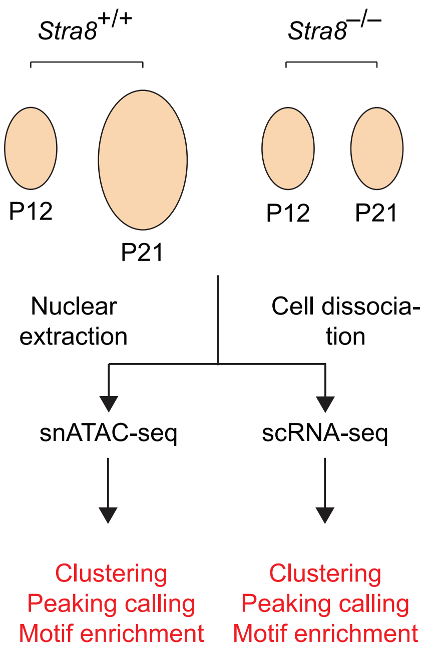

Stra8+/+
Stra8−/−
P12
P21
P12
P21
Nuclear extraction
Cell dissociation
snATAC-seq
scRNA-seq
Clustering
Peaking calling
Motif enrichment
Clustering
Peaking calling
Motif enrichment


**fig. S1**

**fig. S2.** QC filtering plots for the (**A**) WT_P12 data, (**B**) WT_P20 data, (**C**) *Stra8*-KO_P12 data or (**D**) *Stra8*-KO_P20 data from ArchR, showing the transcription start site (TSS) enrichment score and fragment size distributions across individual experiments. (**E**) Violin and box-whisker plot of the number of total aligned fragments for each single cell passing filter per experimental sample and the normalized TSS enrichment for each single cell passing filter per experimental sample. (**F**) Aggregated snATAC-seq fragment size distributions across individual experiments spanning ATAC-seq fragments. Aggregate TSS insertion profiles centered at all TSS regions or fragment size distributions for the cells passing ArchR QC thresholds for each sample in the dataset. Line color represents the sample from the dataset as indicated above the plot.

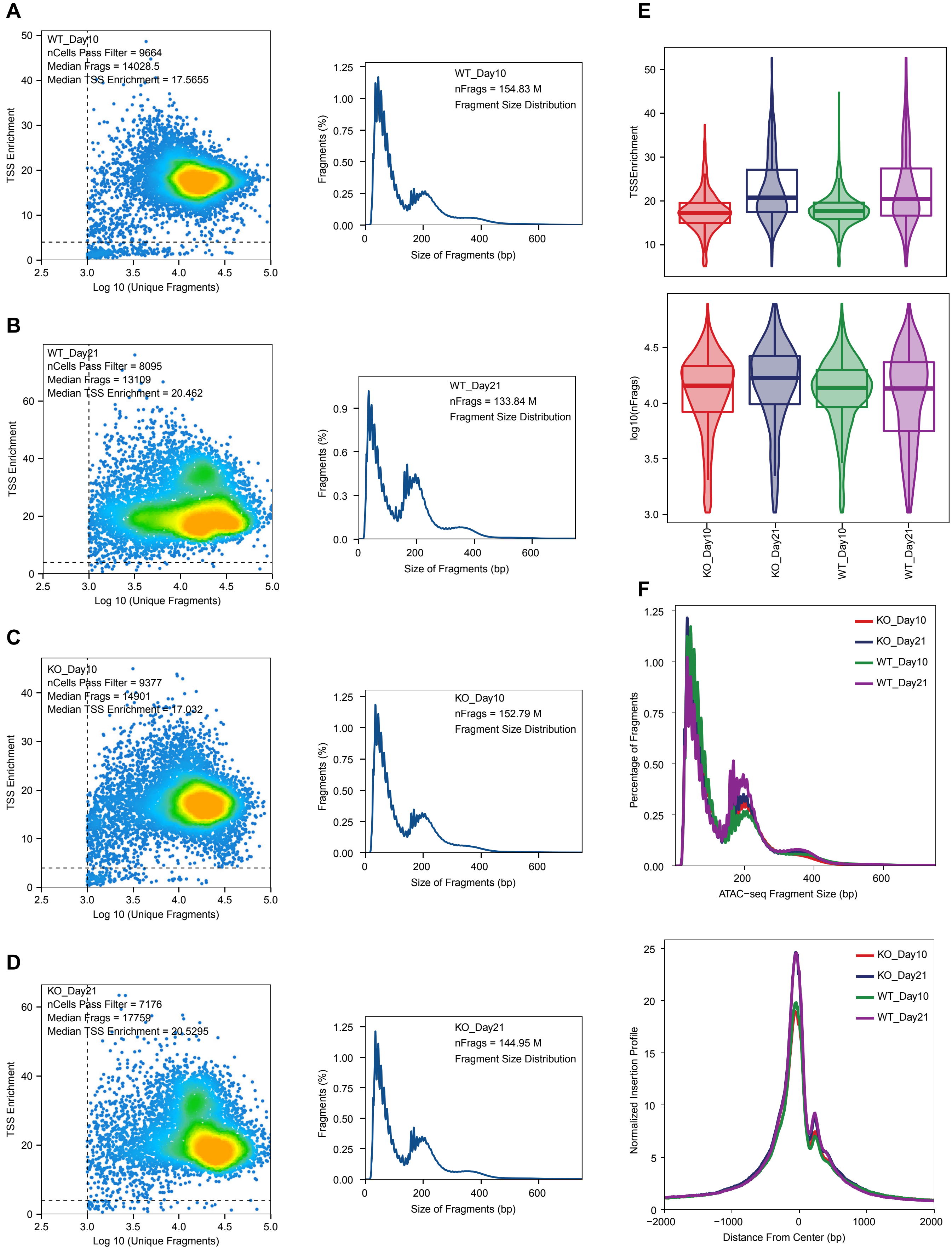

A
WT_Day10
nCells Pass Filter = 9664
Median Frags = 14028.5
Median TSS Enrichment = 17.5655
TSS Enrichment
Log 10 (Unique Fragments)
WT_Day10
nFrags = 154.83 M
Fragment Size Distribution
Fragments (%)
Size of Fragments (bp)
B
WT_Day21
nCells Pass Filter = 8095
Median Frags = 13109
Median TSS Enrichment = 20.462
WT_Day21
nFrags = 133.84 M
Fragment Size Distribution
C
KO_Day10
nCells Pass Filter = 9377
Median Frags = 14901
Median TSS Enrichment = 17.032
KO_Day10
nFrags = 152.79 M
Fragment Size Distribution
D
KO_Day21
nCells Pass Filter = 7176
Median Frags = 17759
Median TSS Enrichment = 20.5295
KO_Day21
nFrags = 144.95 M
Fragment Size Distribution
E
TSSEnrichment
log10(nFrags)
KO_Day10
KO_Day21
WT_Day10
WT_Day21
F
Percentage of Fragments
ATAC−seq Fragment Size (bp)
KO_Day10
KO_Day21
WT_Day10
WT_Day21
Normalized Insertion Profile
Distance From Center (bp)


**fig. S2**

**fig. S3.** (**A-B**) Violin and box-whisker plot of the number of total aligned fragments and normalized transcription start site (TSS) enrichment for each single cell passing filter by WT and *Stra8*-KO sample. Line plot of TSS insertion profiles centered at all TSS regions and fragment size distributions for the cells passing ArchR QC thresholds for WT and *Stra8*-KO sample. (**C-D**) Violin and box-whisker plot of the number of total aligned fragments and normalized transcription start site (TSS) enrichment for each single cell passing filter by P12 and P21 samples. Line plot of TSS insertion profiles centered at all TSS regions and fragment size distributions for the cells passing ArchR QC thresholds for P12 and P21 samples. (**E**) Aggregate violin and box-whisker plot of aligned fragments and normalized transcription start site (TSS) enrichment for the cells passing ArchR QC thresholds for each cell clusters in dataset.
(**F**) Number of cells passing filter for each cell clusters (Unique nuclear fragments > 1,000 and TSS enrichment > 8). (**G**) Aggregate TSS insertion profiles centered at all TSS regions or fragment size distributions for the cells passing ArchR QC thresholds for each cell clusters. Line color represents the sample from the dataset as indicated above the plot.

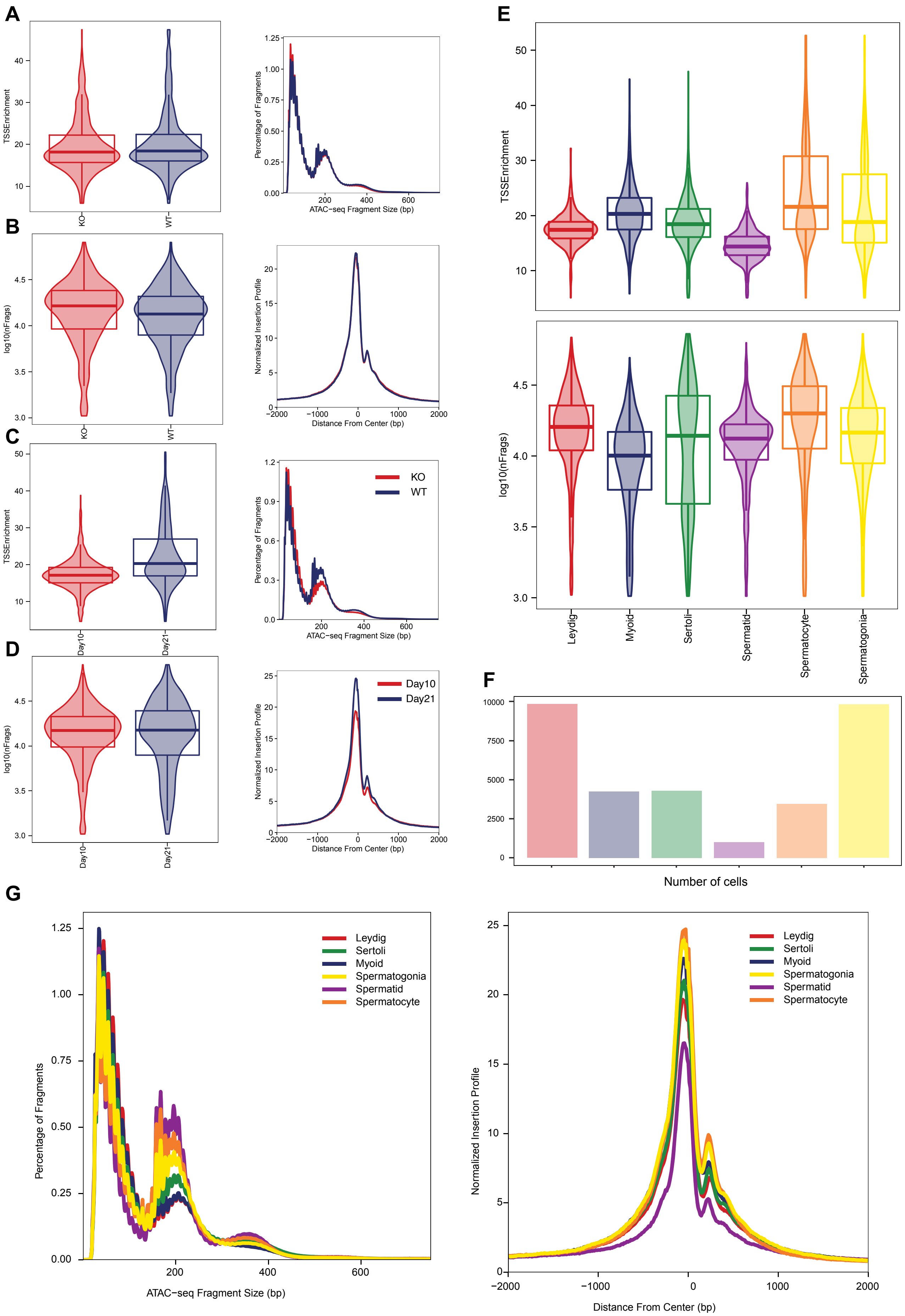
A
TSSEnrichment
KO
WT
Percentage of Fragments
ATAC−seq Fragment Size (bp)
B
log10(nFrags)
Normalized Insertion Profile
Distance From Center (bp)
C
Day10
Day21
KO
WT
D
Day10
Day21
E
Leydig
Myoid
Sertoli
Spermatid
Spermatocyte
Spermatogonia
F
Number of cells
G
Leydig
Sertoli
Myoid
Spermatogonia
Spermatid
Spermatocyte

**fig. S3**

**fig. S4.** (**A**-**D**) QC metrics. (**E**-**F**) Bar plot showing the proportion of the different cell clusters at WT and *Stra8*-KO or P12 and P21. (**G**-**H**) Violin plot showing the X chromosome and Y chromosome ratio of the different cell clusters at WT and *Stra8*-KO or P12 and P21.

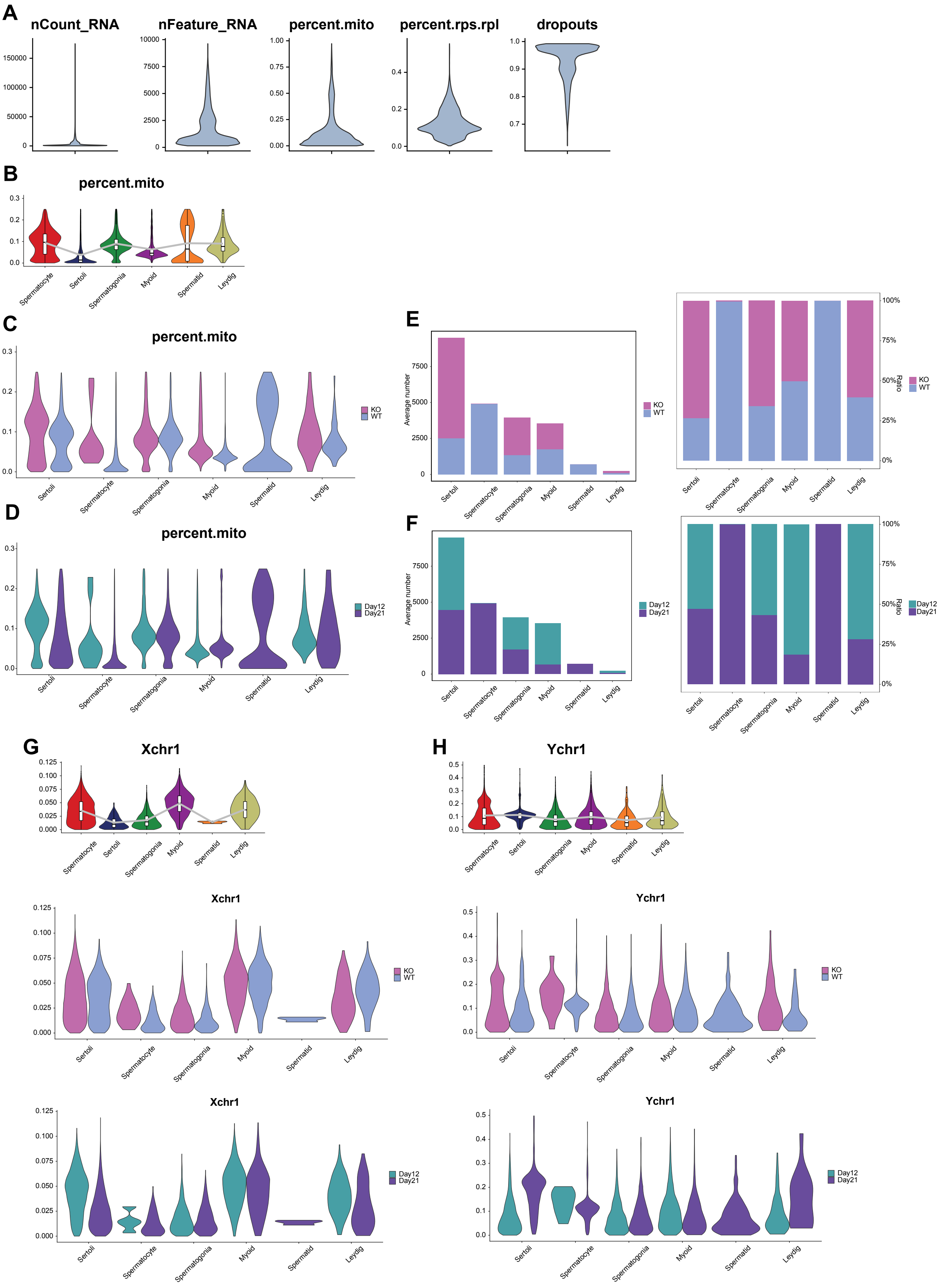
A
nCount_RNA
nFeature_RNA
percent.mito
percent.rps.rpl
dropouts
B
percent.mito
Spermatocyte
Sertoli
Spermatogonia
Myoid
Spermatid
Leydig
C
percent.mito
KO
WT
D
percent.mito
Day12
Day21
E
Average number
Ratio
F
G
Xchr1
H
Ychr1

fig. S4

**fig. S5.** (**A**) A UMAP plot for analyzed cell cluster and samples from 32,657 cells in snATAC-seq dataset. (**B**) A bar plot showing the proportion of the different cell clusters at different time points in scATAC-seq. (**C**) Gene-activity scores for *Stra8*, *Syce1*, *Acrv1*, *Sox9*, *Acta2*, and *Star*. (**D**) Genome accessibility track visualization of *Stra8*, *Syce1*, *Acrv1*, *Sox9*, *Acta2*, and *Star*.

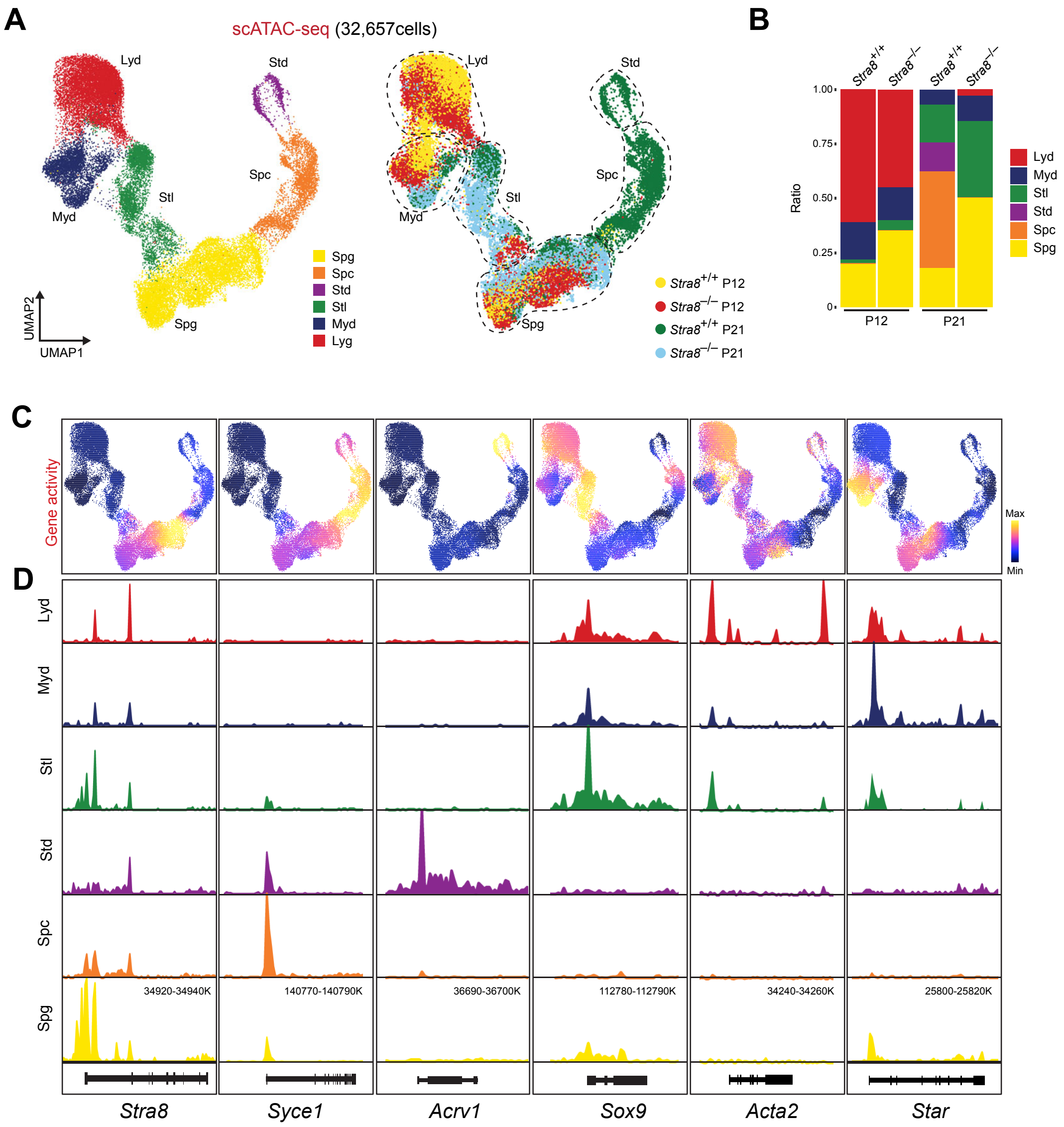
A
scATAC-seq (32,657cells)
Lyd
Std
Spc
Myd
Stl
Spg
UMAP2
UMAP1
Spg
Spc
Std
Stl
Myd
Lyg
Strain8+/+ P12
Stra8−/− P12
Stra8+/+ P21
Stra8−/− P21
B
Stra8+/+
Stra8−/−
Ratio
1.00
0.75
0.50
0.25
0
P12
P21
Lyd
Myd
Stl
Std
Spc
Spg
C
Gene activity
Max
Min
D
Lyd
Myd
Stl
Std
Spc
Spg
34920-34940K
140770-140790K
36690-36700K
112780-112790K
34240-34260K
25800-25820K
Stra8
Syce1
Acrv1
Sox9
Acta2
Star

**fig. S5**

**fig. S6.** (**A**) A UMAP plot for analyzed cell cluster and samples from 22,839 cells in scRNA-seq dataset. (**B**) A bar plot showing the proportion of the different cell clusters at different time points in scRNA-seq. (**C**) Gene expression for *Stra8*, *Syce1*, *Acrv1*, *Sox9*, *Acta2*, and *Star*. (**D**) Violin plots showing the expression level of marker genes in each cluster. (**E**) Heatmap showing Pearson's correlation coefficients between snATAC-seq gene activity scores and gene expression values in dataset. Each column represents a cell type in the scRNA-seq data and each row represents a cell type in the scATAC-seq data.

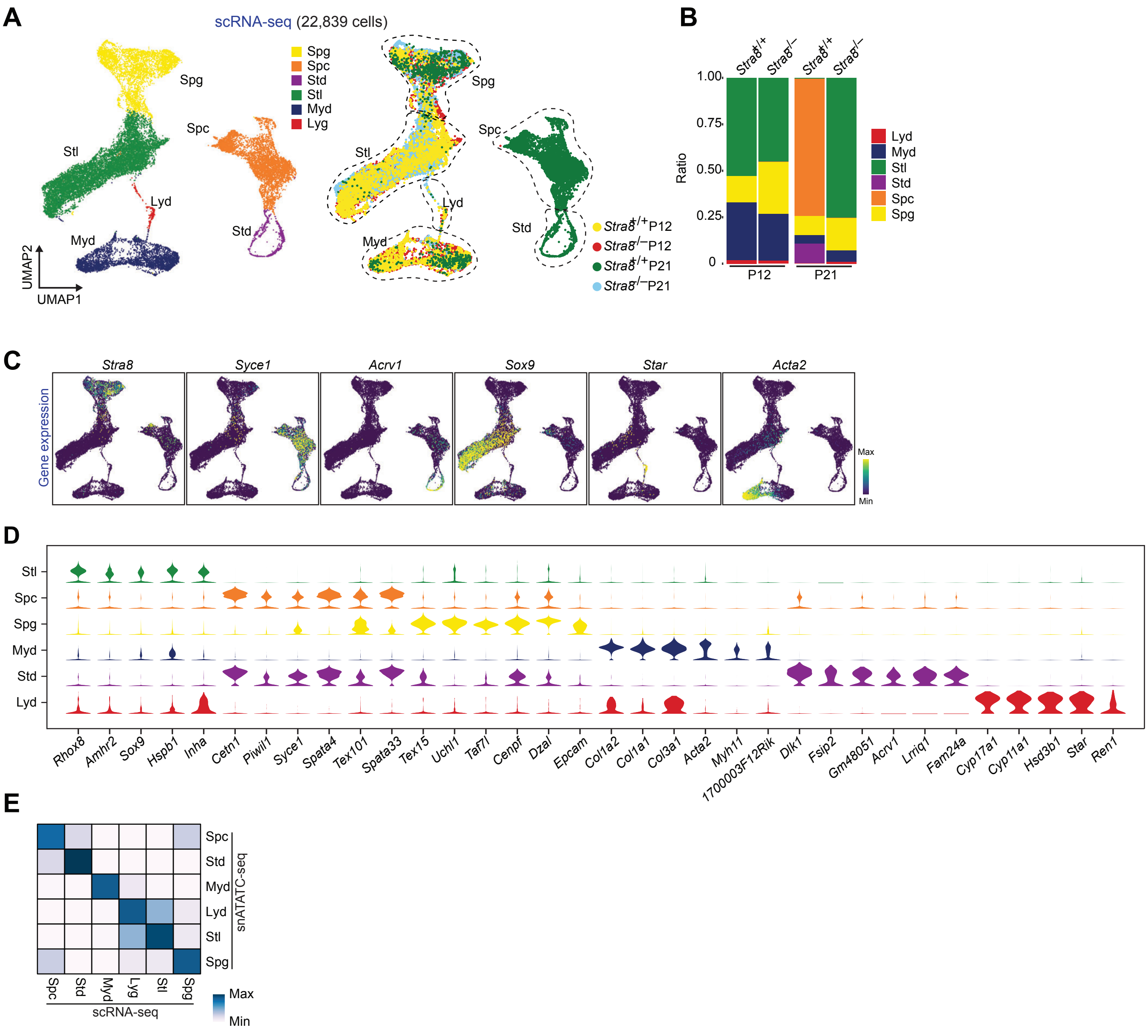
A
scRNA-seq (22,839 cells)
Spg
Spc
Stl
Lyd
Myd
Std
Spg
Spc
Std
Stl
Myd
Lyg
UMAP2
UMAP1
Strain8+/+ P12
Strain8-/- P12
Strain8+/+ P21
Strain8-/- P21
B
Ratio
1.00
0.75
0.50
0.25
0
P12
P21
Lyd
Myd
Stl
Std
Spc
Spg
C
Gene expression
Stra8
Syce1
Acrv1
Sox9
Star
Acta2
Max
Min
D
Stl
Spc
Spg
Myd
Std
Lyd
Rhox8
Amhr2
Sox9
Hspb1
Inha
Cetn1
Piwil1
Syce1
Spata4
Tex101
Spata33
Tex15
Uchl1
Taf7l
Cenpf
Dzal
Epcam
Col1a2
Col1a1
Col3a1
Acta2
Myh11
1700003F12Rik
Dlk1
Fsip2
Gm48051
Acrv1
Lrriq1
Fam24a
Cyp17a1
Cyp11a1
Hsd3b1
Star
Ren1
E
Spc
Std
Myd
Lyd
Stl
Spg
snATATC-seq
Spc
Std
Myd
Lyg
Stl
Spg
scRNA-seq
Max
Min

**fig. S6**
